# G3BP1 Maintains Lysosomal Homeostasis to Limit Tau Aggregate Accumulation

**DOI:** 10.1101/2025.08.29.672259

**Authors:** Tomo Kimura, Riku Saito, Mariko Okada, Satoaki Matoba, Atsushi Hoshino, Yoshihisa Watanabe

## Abstract

Tau aggregation is a pathological hallmark of a group of neurodegenerative diseases collectively termed tauopathies. While impaired proteostasis is known to drive the accumulation of abnormal proteins, the molecular factors influencing Tau aggregate clearance remain incompletely understood. In this study, we employed a cell-based Tau reporter assay combined with a lentivirus-based pooled CRISPR-Cas9 loss-of-function library to identify genes that regulate Tau aggregation. Genome-wide screening revealed that candidate genes were significantly enriched in categories related to mRNA metabolic processes and autophagy. Among them, we focused on the RNA-binding protein G3BP1, which is functionally associated with both processes. Detailed analyses showed that G3BP1 deficiency promoted the accumulation of Tau aggregates without affecting stress granule formation or autophagic flux. Instead, G3BP1 dysfunction resulted in impaired lysosomal homeostasis, as evidenced by reduced lysosomal abundance and acidification. Furthermore, lysosomal damage induced by L-leucyl-L-leucine methyl ester (LLOMe) enhanced Tau aggregation, particularly in G3BP1-deficient cells. Conversely, pharmacological activation of TFEB by the curcumin analog C1 restored lysosomal function and suppressed Tau aggregate accumulation to wild-type levels. These findings highlight a role of G3BP1 in maintaining lysosomal homeostasis and promoting Tau clearance. Our results further suggest that therapeutic strategies aimed at enhancing lysosomal biogenesis, such as TFEB activation, may hold promise for the treatment of tauopathies.

## 1 INTRODUCTION

Tau (MAPT) regulates microtubule assembly and stabilizes microtubule networks (Parra Bravo *et al*. 2024). The abnormal accumulation of Tau leads to various neurodegenerative diseases, including Alzheimer’s disease (AD), frontotemporal dementia (FTD), progressive supranuclear palsy (PSP), and corticobasal degeneration (CBD) (Samudra *et al*. 2023). In the brains of patients with these diseases, filamentous and hyperphosphorylated Tau accumulates as a component of neurofibrillary tangles (NFTs) and is a pathological hallmark of tauopathies (Parra Bravo *et al*. 2024). There are six isoforms of Tau generated by alternative splicing (Goedert *et al*. 1992). In particular, isoforms with different numbers of repeat domains in the microtubule-binding region, known as 3R and 4R Tau, are also a focus of neuropathological interest. Both 3R and 4R Tau isoforms are found in NFTs in the brains of patients with AD, chronic traumatic encephalopathy, and primary age-related tauopathy (Zhang *et al*. 2022). Tau deposits in PSP and CBD are predominantly composed of 4R Tau, whereas deposits in Pick’s disease primarily contain 3R Tau (Zhang *et al*. 2022).

Pathological proteins in various neurodegenerative diseases, such as Tau, abnormally aggregate and accumulate in the brain (Goedert *et al*. 1992). In AD, amyloid-β, along with Tau, accumulates as aggregates; in Parkinson’s disease (PD), α-synuclein forms aggregates; and in amyotrophic lateral sclerosis (ALS), proteins such as TDP-43 and FUS aggregate (Dugger & Dickson 2017; Portz *et al*. 2021). The accumulation of abnormal proteins exerts cytotoxic effects, leading to neuronal degeneration. In healthy individuals, proteostasis mechanisms, such as molecular chaperone-mediated protein folding and degradation via autophagy and the proteasome, prevent the accumulation of abnormal proteins (Hipp *et al*. 2014). However, aging, unhealthy lifestyle, and genetic factors cause proteostasis imbalance, resulting in various neurodegenerative diseases (Hipp *et al*. 2014).

There is substantial evidence indicating that protein aggregates associated with neurodegenerative diseases can be transmitted between brain cells, contributing to the spread of pathology (Stopschinski & Diamond 2017). The brain-derived or *in vitro*–generated fibrils of α-synuclein and Tau act as seeds and promote the aggregation of their soluble forms in the rodent brain and cultured cells (Luk *et al*. 2012; Sanders *et al*. 2014; Volpicelli-Daley *et al*. 2011; Frey *et al*. 2023). Such experimental systems have proven to be highly valuable for elucidating the mechanisms of disease progression and for providing tools to explore potential therapeutic interventions. In this study, we employed a cell-based Tau aggregate reporter assay as described above to identify factors that influence aggregate accumulation. Using a lentivirus-based pooled CRISPR library as a comprehensive genetic screening tool, we identified multiple candidate genes, including those involved in mRNA metabolic processes and autophagy. Given the close association of G3BP1 with both mRNA metabolism and autophagy, we focused on this factor to explore its role in Tau aggregate clearance. Our findings indicate that loss of G3BP1 disrupts lysosomal homeostasis and promotes the accumulation of Tau aggregates. Moreover, activation of TFEB by the curcumin analog C1 was able to suppress Tau aggregate accumulation in G3BP1-deficient cells.

## 2. Material & Methods

### 2.1 Cell Culture and lentivirus production

HEK293 or HEK293T cells were grown in DMEM medium (Nacalai Tesque, Kyoto, Japan) supplemented with 10% fetal bovine serum (Sigma-Aldrich, St. Louis, MO,) and 1% penicillin-streptomycin mixed solution (Nacalai Tesque). Cells were transfected with plasmids (pAAV-mCherry, pAAV-G3BP1, and pAAV-GFP) using FuGENE HD (Promega, Madison, WI, USA). Lentiviral particles were produced by co-transfecting HEK293T cells with the expression plasmids (pLenti-TauRD-GFP, pLenti-TauFL-K294I/P301L and pLenti-TFEB-mNeoGreen(mNG)), the packaging plasmid psPAX2, and the envelope plasmid pMD2.G. After transfection for 24 h, the medium was changed by DMEM medium containing 10% FBS. The supernatants were collected after transfection for 48 h and were subjected to centrifugation at 5,000 rpm for 5 min to remove cell precipitate. Cells were infected by adding the viral supernatant to the culture medium. After two days, the medium was replaced, and the cells were further cultured prior to analysis. To induce stress granule formation, cells were subjected to oxidative stress by treatment with 200 μM sodium arsenite (Merck Millipore, Burlington, MA) for 1 h (Mackenzie *et al*. 2017).

### 2.2 Tau aggregation assay

The Tau aggregation assay was performed according to the method developed by Stöhr et al (Stöhr *et al*. 2017). TauRD-GFP aggregation was assessed in cells expressing a reporter protein consisting of the four microtubule-binding repeats of Tau (residues 244–372) containing the P301L mutation fused to GFP. To promote aggregation, PFF seeds generated from a 31-amino acid peptide (residues 306–336; AnaSpec, Fremont, CA) within the third repeat domain of Tau were introduced into the cells by lipofection. Two days after seed introduction, the cells were dissociated with trypsin, collected, and resuspended in FACS buffer containing 0.1% saponin. Saponin treatment allows soluble TauRD-GFP to diffuse out of the cells, whereas aggregates remain intracellular. The number of TauRD-GFP aggregate–positive cells was then determined by Attune NxT Flow Cytometer (Thermo Fisher Scientific, Waltham, MA). For the full-length Tau aggregation assay, a construct was generated in which GFP was fused to Tau carrying the aggregation-prone mutations K294I and P301L (Tau^K294I/P301L^). Tau^K294I/P301L^-GFP was transiently expressed in HEK293 cells, and aggregate-positive cells were measured by flow cytometry using the same procedure as described above.

### 2.3 CRISPR Library Screening

Stable Cas9 endonuclease expression was established in TauRD-GFP–expressing HEK293 cells prior to sgRNA lentiviral library transduction. To prevent multiple viral transduction events, cells were infected with the pooled sgRNA lentivirus at a low multiplicity of infection (MOI). Following genome editing in individual cells, Tau PFF seeds were introduced to promote TauRD-GFP aggregation. At two days post–PFF seed introduction, cells were collected and subjected to membrane permeabilization with 0.1% saponin. Cells were then sorted into high- and low-aggregate populations using a Cell Sorter MA900 (Sony Biotechnology, Tokyo, Japan). Genomic DNA was prepared from the sorted cells, and the integrated sgRNA regions were amplified by PCR to identify the sgRNA sequences. Next-generation sequencing was performed on the pooled amplicons to determine their nucleotide sequences, and sgRNAs from sorting cells were counted.

### 2.4 Enrichment Analysis

Analysis of deep sequencing reads was performed using MAGeCK analysis (Li *et al*. 2014). Gene enrichment analysis was then conducted to identify candidate genes associated with Tau aggregation. The enrichment of sgRNAs between GFP-high and GFP-low populations was compared, and scatter plots were generated using ggVolcanoR v1.0 (Mullan *et al*. 2021). Gene Ontology (GO) enrichment analysis of candidate genes was carried out with the Metascape bioinformatics tool (Zhou *et al*. 2019).

### 2.5 G3BP1 and G3BP2 Knockout using CRISPR/Cas9

To generate G3BP1-, G3BP2-, or G3BP1/2-knockout (KO) cells, TrueGuide synthetic gRNAs targeting human G3BP1 or G3BP2 were purchased from Thermo Fisher. gRNAs, TrueCut Cas9 Protein v2 (Thermo Fisher) and Lipofectamine Cas9 Plus Reagent (Thermo Fisher) were mixed into OPTI-MEM (Thermo Fisher). Diluted Lipofectamine CRISPRMAX Reagent (Thermo Fisher) was added to the mixture, and after a 15 min incubation, the resulting complex was delivered to HEK293 cells. KO cells were isolated by limiting dilution. The knockout was verified by PCR amplification of genomic DNA flanking the target site, followed by cloning into a plasmid vector and DNA sequencing to confirm the knockout at the genomic level. Sequencing revealed that each cloned cell harbored 326-bp insertion, 124-bp insertion and 2-bp deletion in exon 4 of G3BP1, and 198-bp insertion, 4-bp deletion and 18-bp deletion in exon 4 of G3BP2. Furthermore, western blotting of G3BP1 and G3BP2 confirmed that each protein was absent in the respective knockout cells (Figure S1).

### 2.6 Western Blotting

For western blotting, cell lysates were resolved on 10% SDS-PAGE gels (Fuji Film, Osaka, Japan) and subsequently transferred to PVDF membranes (Merck Millipore). PVDF membranes were blocked with 10% skim milk and incubated with anti-G3BP1 (1:1000 dilution; Abcam, Cambridge, UK), anti-G3BP2 (1:1000 dilution; Cell Signaling Technology, Danvers, MA) and anti-Actin antibody (1:2000 dilution; ProteinTech, Rosemont, IL) for 24 h. Following membrane washing, an alkaline phosphatase–conjugated secondary antibody (Jackson ImmunoResearch Laboratories, West Grove, PA) was applied. Immunopositive signals were detected with nitroblue tetrazolium chloride (Nacalai Tesque) and 5-bromo-4-chloro-3-indolylphosphate P-toluidine salt (Nacalai Tesque) reagents. Western blot images were scanned and converted to digital format using an EPSON GT-X970 scanner (Seiko Epson, Suwa, Japan).

### 2.7 Immunocytochemistry

Immunocytochemical analysis was performed as described previously (Watanabe *et al*. 2017). Cells were fixed with 4% paraformaldehyde (Merck Millipore) for 30 min. The fixed cells were permeabilized in PBS containing 0.1% Triton X-100 and placed in 5% normal goat serum for 30 min. Cells were incubated with anti-LAMP-1 (1:1000 dilution; Santa Cruz Biotechnology, Santa Cruz, CA) or anti-TIA-1 (1:1000 dilution; ProteinTech) antibody for 12 h at 25°C. After a PBS wash, cells were further incubated with Alexa Fluor Plus 488-labeled goat anti-mouse IgG or Alexa Fluor Plus 488-labeled goat anti-rabbit IgG (Thermo Fisher) for 4 h at 25°C. After washing, nuclei were stained with 4′,6-diamidino-2-phenylindole (Dojindo, Kumamoto, Japan). The cells on cover glasses were mounted on glass slides in an aqueous mounting medium (FluorSave Reagent; Merck Millipore) and imaged using a confocal laser microscope (LSM900; Zeiss, Oberkochen, Germany).

### 2.8 Autophagy Flux analysis

Autophagic flux analysis was performed using the GFP-LC3-RFP probe (Kaizuka *et al*. 2016). pMRX-IP-GFP-LC3-RFP plasmid was transfected into cells. Cells were cultured in puromycin-containing medium for one week to select stable expression clones. Following trypsinization, cells were washed with FACS buffer and subjected to flow cytometric analysis to measure GFP and RFP fluorescence intensities. Relative autophagic flux was calculated as the RFP/GFP fluorescence ratio.

### 2.9 Measurement of lysosomal activity

For the measurement of lysosomal abundance and acidification, LysoTracker Red DND-99 (Thermo Fisher), LysoPrime Deep Red (Dojindo), and pHLys Green (Dojindo) were used. LysoTracker Red DND-99 was diluted in serum-reduced culture medium and applied to the cells for 2 h to allow uptake. LysoPrime Deep Red was diluted and applied to the cells for 30 min, followed by incubation with pHLys Green for 2 h. After trypsinization, cells were washed with FACS buffer and analyzed by flow cytometry to measure the fluorescence intensity of each dye.

### 2.10 Measurement of TFEB activity

TFEB-mNG was introduced into WT and G3BP1 KO cells using a lentiviral vector, and cells were treated with either 1 μM curcumin analog C1 or 10 μM rapamycin for 24 h. After fixation with 4% PFA, nuclei were stained with PBS containing 0.1% Triton X-100 and DAPI, and images were acquired using confocal laser scanning microscopy. Nuclear and total cellular mNG fluorescence intensities were quantified using ImageJ, and relative TFEB activity was expressed as the ratio of nuclear to total cellular TFEB-mNG fluorescence intensity. A total of 76 cells per condition were analyzed and plotted. Statistical significance was determined by two-way ANOVA followed by Tukey’s multiple-comparison test.

### 2.11 Lysosomal damage and enhancement of lysosomal biogenesis

Lysosomal damage was induced by treatment with LLOMe (Cayman Chemical, Ann Arbor, MI) (Maejima *et al*. 2013). For monitoring lysosomal damage, cells were transfected with a DsRed-Galectin-3 reporter, and its subcellular localization was examined using fluorescence microscopy (IX70; Evident, Tokyo, Japan). For the TauRD-GFP aggregation assay, LLOMe was added immediately after PFF seed introduction, and aggregates were quantified two days later. Lysosomal biogenesis was promoted using curcumin analog C1 (TFEB activator 1; MedChemExpress, Monmouth Junction, NJ). To evaluate the activation of lysosomes by 1μM curcumin analog C1, lysosomal activity was measured using the aforementioned lysosomal fluorescent dyes. Cells were pretreated with curcumin analog C1 for 24 h prior to PFF seed introduction, and subsequently cultured for 48 h in medium containing TFEB activator 1. TauRD-GFP aggregation was then assessed as described above.

### 2.12 Statistics and reproducibility

GraphPad Prism software was used for statistical analysis. All experiments were performed in triplicate, and statistical values were displayed as the means□±□SD (standard deviation). *p*□>□0.05 was considered not significant. \**p* < 0.05, \**p*□<L0.01, \*\*\**p*□<L0.001 by Student’s t-test and one-way ANOVA. Tukey’s *post hoc* test was performed.

## 3 RESULTS

### 3.1 Genome-Wide CRISPR Screening for Tau Aggregation

Currently, CRISPR-Cas9 libraries are widely used as powerful tools for analyzing gene function. We conducted pooled CRISPR-Cas9 loss-of-function screens to elucidate genes associated with Tau aggregate accumulation. We utilized HEK293 cells expressing a reporter in which GFP was fused to the Tau repeat domain (Tau244–372) carrying the P301L mutation (TauRD-GFP) to monitor Tau aggregates. When preformed fibrils (PFFs) prepared from the third repeat domain peptide (R3, residues 306–336) were introduced into these reporter cells, aggregation of the reporter protein was promoted (Stöhr *et al*. 2017). Initially, the reporter cells were stably transduced with Cas9 and subsequently infected at a low MOI with a pooled lentiviral sgRNA library, ensuring that each cell expressed a unique sgRNA (Figure 1A). Subsequently, PFF seeds were delivered into the cells by lipofection to induce aggregation of TauRD-GFP. To specifically detect TauRD-GFP aggregates, the cells were treated with saponin to permeabilize the plasma membrane, allowing the soluble form to be released into the extracellular space. The cells were subjected to FACS-based screening and sorted into populations with high and low levels of aggregation (Figure 1 A). To identify the sgRNAs present in each cell population, genomic DNA was collected, and guide sequences amplified by PCR were then identified by deep sequencing. We compared sgRNA counts between the high- and low-aggregation populations, and visualized the genes enriched in the high-aggregation group in a scatter plot (Figure 1B, Table S1, S2). Metascape enrichment GO analysis of genes that significantly influenced TauRD-GFP aggregate accumulation revealed that disruptions in mRNA metabolic processes (GO:0016071), autophagy (GO:0006914), and DNA metabolic processes (GO:0051052) contribute to this accumulation (Figure 1C, D). Previous studies have clarified that autophagy has a strong impact on the accumulation of protein aggregates in various neurodegenerative diseases, including Alzheimer’s disease and Parkinson’s disease (Arotcarena *et al*. 2019; Angst *et al*. 2025; Watanabe *et al*. 2012). Consistent with this notion, our present analysis revealed multiple autophagy-related genes (40 candidate genes), such as ATG7 and ATG16L1, as strong candidate hits (Figure1B, Table S3). In the mRNA metabolic process, 66 candidate genes, including DHX8, HNRNPF, and SNRNP25, were identified (Table S3).

**Figure 1.**
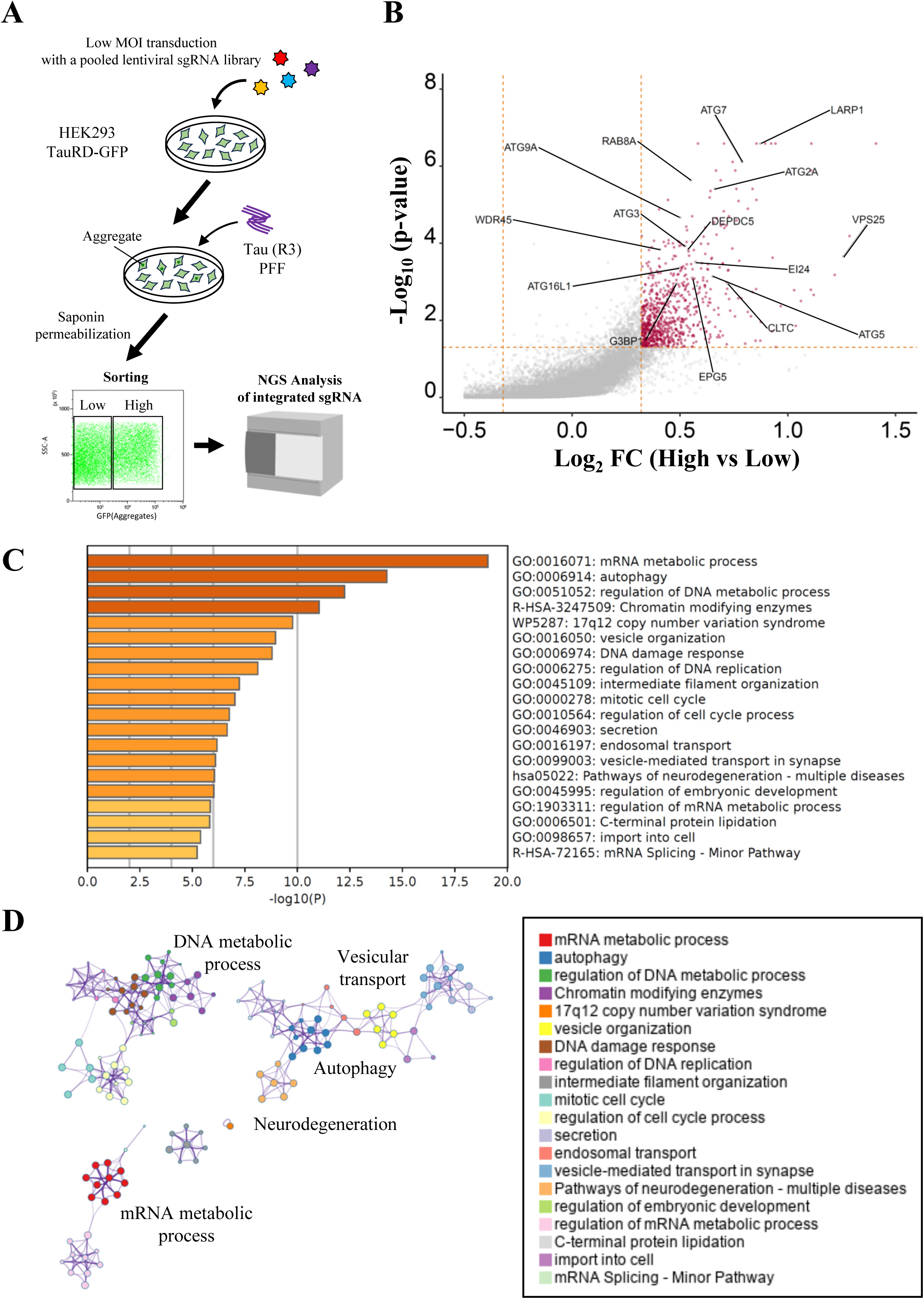
CRISPR library screening for Tau pathology using a cell-based assay and enrichment analysis. **A**. Schematic workflow of the CRISPR-Cas9 library screening for Tau pathology is shown. HEK293 reporter cells stably expressing Cas9 and TauRD-GFP were transduced with a pooled sgRNA library, seeded with PFFs, and sorted into high- and low-aggregation populations by FACS. Genomic DNA was extracted from each population, and sgRNA sequences were amplified and identified by deep sequencing. **B**. Candidate genes enriched in the high-aggregation population were analyzed statistically and plotted on a scatter plot. *n* = 4, *p* < 0.05, Fold changes (FC) >1.25. **C**. Candidate genes were subjected to GO enrichment analysis, and significantly overrepresented GO terms were highlighted. **D**. Furthermore, functional networks generated by clustering the results of the GO enrichment analysis are depicted.

### 3.2 Deficiency of G3BP1 Leads to Increased Accumulation of Tau Aggregates

Among these genes, we focused on G3BP1 and investigated in detail the mechanisms underlying the increased accumulation of Tau aggregates. Then, G3BP1 KO cells were generated, and the accumulation of TauRD-GFP reporter aggregates was measured (Figure 2A, 2B). In the absence of PFF-induced aggregation, the TauRD-GFP reporter did not form aggregates even in G3BP1 KO cells. By contrast, there was a significant accumulation of PFF-induced TauRD-GFP aggregates in G3BP1 KO cells, consistent with the findings from the CRISPR-Cas9 library screen. Furthermore, exogenous expression of G3BP1 in the KO cells reduced the accumulation of PFF-induced TauRD-GFP aggregates (Figure 2B, Figure S2A), ruling out the possibility that this phenotype resulted from CRISPR/Cas9 off-target effects. We next assessed aggregate accumulation in full-length Tau containing aggregation-prone mutations (Tau^K294I/P301L^-GFP). When transiently overexpressed, this Tau mutant forms aggregates exhibits AT8 immunoreactivity in cells even in the absence of PFF seed introduction (Figure 2C, 2D). The proportion of aggregate-positive cells was significantly higher in G3BP1 KO cells than in WT cells, as shown in Figure 2D.

**Figure 2.**
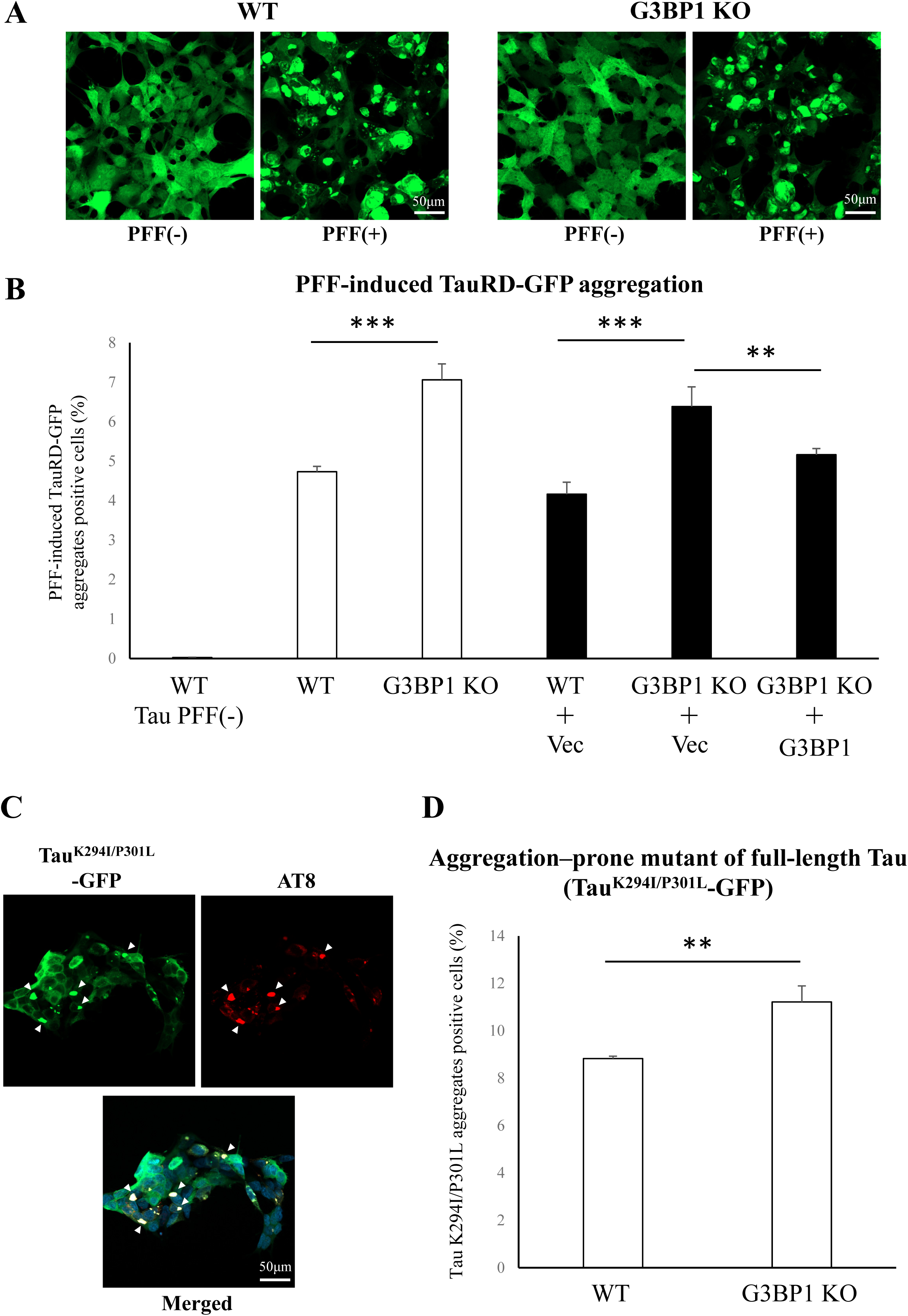
Increased accumulation of Tau aggregates in G3BP1 KO cells. **A.** Tau R3 PFF seed induces TauRD-GFP aggregation. **B**. Accumulation of PFF seed–induced TauRD-GFP aggregates was compared between WT and G3BP1 KO cells. WT and G3BP1 KO cells (*open bars*). To rule out off-target effects, the same analysis was performed after exogenous introduction of either a mock vector or a G3BP1 expression vector (*closed bars*). The data are expressed as means ± SD (*n* = 3). Statistical analysis was performed using one-way ANOVA followed by Tukey’s *post-hoc* test. *p\*\** < 0.01 and *p\*\*\** < 0.001. **C**. Aggregation–prone mutant of full-length Tau (Tau^K294I/P301L^-GFP) accumulated as large AT8-positive aggregates. **D**. Tau^K294I/P301L^-GFP is expressed in WT and G3BP1 KO cells, and aggregate-positive cells were quantified by flow cytometry. The data are expressed as means ± SD (*n* = 3). Statistical analysis was performed using Student’s *t*-test. *p\*\** < 0.01.

### 3.3 Stress Granule Formation and Autophagic Activity in G3BP1 KO Cells

G3BP1 is known to possess functions related to stress granule formation. When cells are exposed to various stresses, such as heat shock or oxidative stress, translation is temporarily halted to protect cellular functions (Van Treeck & Parker 2019). During this period, granular structures containing RNA form in the cytoplasm, which disappear once the stress is alleviated. Stress granules are closely linked to neurodegenerative diseases, and ALS-associated proteins, including TDP-43 and FUS, have been shown to localize to these granules (Wolozin & Ivanov 2019). To clarify how G3BP1 loss leads to cellular dysfunction and Tau aggregate accumulation, we assessed stress granule formation in G3BP1 KO cells. Cells were exposed to sodium arsenite for 1 h, after which the localization of the stress granule marker TIA-1 was assessed by immunostaining. G3BP1 KO cells exhibited normal stress granule formation comparable to that of wild-type (WT) cells; however, G3BP1/2 double KO cells were unable to form stress granules (Figure S3A). Next, we measured autophagic activity in G3BP1 KO cells. Autophagic activity in WT and KO cells stably expressing the GFP-LC3-RFP probe was measured by FACS, and no significant difference was observed (Figure S3B). These results indicate that the accumulation of Tau aggregates in G3BP1-deficient cells may not be attributable to defects in stress granule formation or autophagic function.

### 3.4 Reduced Lysosomal Function in Cells Lacking G3BP1

Next, we examined lysosomal morphology and activity. Immunostaining for the lysosomal marker LAMP-1 revealed no abnormalities in lysosomal morphology or localization in G3BP1 KO cells (Figure 3A). In contrast, FACS analysis of lysosomal uptake of LysoTracker Red DND-99, a cell-permeable red fluorescent dye that stains acidic compartments within a cell, revealed a significant decrease in G3BP1 KO cells compared with WT cells, and exogenous expression of G3BP1 restored the uptake to WT levels. (Figure 3B, Figure S2B). Furthermore, we separately measured lysosomal abundance and acidification using a different fluorescent probe. LysoPrime Deep Red dye and pHLys Green dye are specifically incorporated into lysosomes in a pH-independent and pH-dependent manner, respectively. Cells were simultaneously incubated with both fluorescent dyes, washed, and the uptake of each dye was analyzed by FACS. Compared with WT cells, G3BP1 KO cells showed a significant reduction in the uptake of both LysoPrime Deep Red dye and pHLys Green dye (Figure 3C, D), indicating that G3BP1 deficiency affects both lysosomal abundance and acidification.

**Figure 3.**
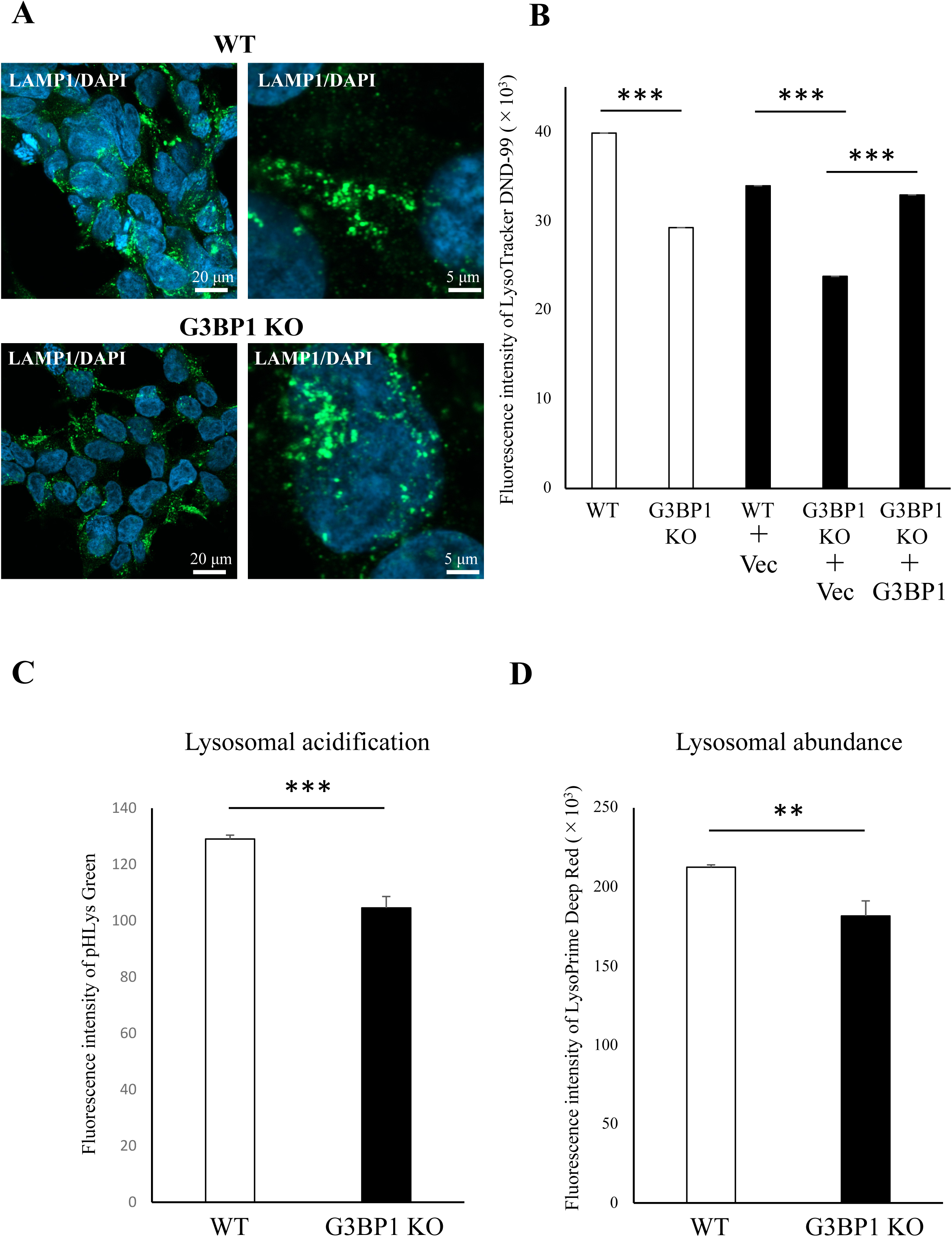
Impaired lysosomal function caused by G3BP1 deficiency. **A**. For morphological analysis of lysosomes, immunostaining of the lysosomal marker LAMP-1 was performed, and the cells were analyzed using confocal laser microscopy (*left panels*). Merged images with DAPI staining are shown (*right panels*). The upper panels show WT cells, and the lower panels show G3BP1 KO cells. **B**. Uptake of LysoTracker Red DND-99 was measured by flow cytometry in WT and G3BP1 KO cells (open bars). In addition, the same experiment was performed after transduction with either a mock vector or a G3BP1 expression vector (closed bars). The data are expressed as means ± SD (*n* = 3). Statistical analysis was performed using one-way ANOVA followed by Tukey’s *post-hoc* test. *p\*\*\** < 0.001. **C**. Lysosomal acidification was assessed by measuring the uptake of the fluorescent dye pHLys Green in WT (open bars) and G3BP1 KO (closed bars) cells. Fluorescence intensity of each cell was quantified by flow cytometry, and the data are presented as means ± SD (*n* = 3). Statistical analysis was performed using Student’s *t*-test. *p\*\*\** < 0.001. **D**. Similarly, lysosomal abundance was measured in each cell type using LysoPrime Deep Red. The data are presented as means ± SD (*n* = 3). Statistical analysis was performed using Student’s *t*-test. *p\*\** < 0.01.

### 3.5 Impact of Lysosomal Activity on Tau Aggregate Accumulation

These findings indicate that G3BP1 may affect the maintenance of lysosomal homeostasis. Therefore, we assessed the impact of lysosomal damage on Tau aggregate accumulation by treating cells with lysosome-disrupting agents. L-Leucyl-L-Leucine methyl ester (LLOMe) is a lysosomotropic compound that, once inside lysosomes, is converted into oligomers by lysosomal cathepsin C, which disrupt the membrane (Uchimoto *et al*. 1999). PFF seeds were introduced into WT or G3BP1 KO cells expressing TauRD-GFP simultaneously with the addition of LLOMe to artificially induce lysosomal damage. Lysosomal damage was confirmed by monitoring with DsRed-Galectin3 (Figure 4A). Treatment with LLOMe led to a significant increase in TauRD-GFP aggregates in both cell types (Figure 4B). This result suggests that severe lysosomal damage leads to massive accumulation of Tau aggregates. Notably, compared with WT cells, the effect of LLOMe-induced lysosomal damage on Tau aggregate accumulation was more pronounced in G3BP1 KO cells (Figure 4B).

**Figure 4.**
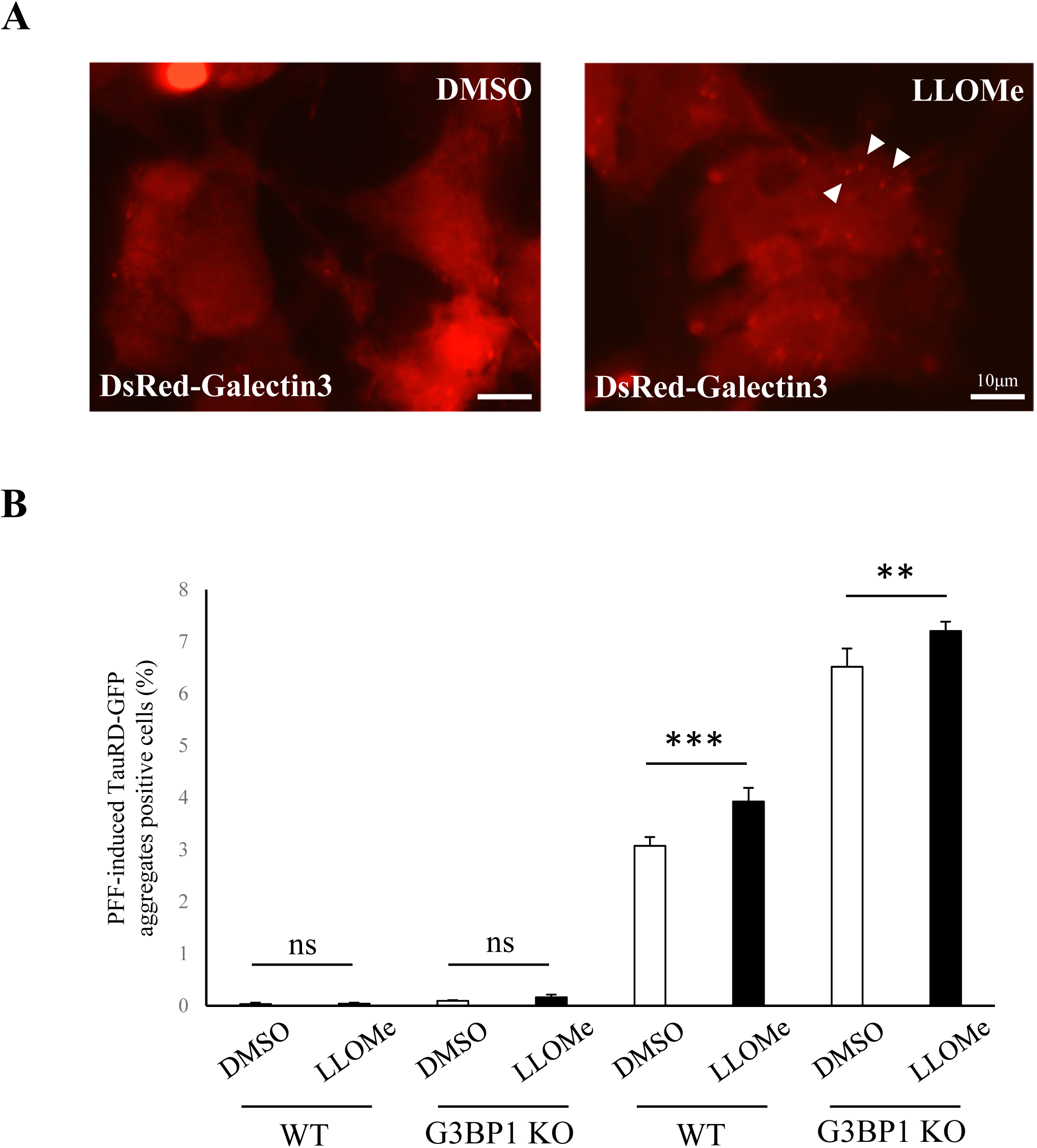
LLOMe-mediated lysosomal damage enhances accumulation of PFF seed-induced Tau aggregates. **A**. Cells transiently expressing DsRed-Galectin3 were treated with DMSO (*upper*) or LLOMe (*lower*), and the subcellular localization of DsRed-galectin3 was visualized by fluorescence microscopy. Arrowheads indicate DsRed-galectin3 puncta, which represent damaged lysosomes. **B**. WT and G3BP1 KO cells were subjected to lysosomal damage under this condition, and PFF seed-induced TauRD-GFP aggregation was assessed. Open bars indicate DMSO-treated cells, while closed bars indicate LLOMe-treated cells. The data are expressed as means ± SD (*n* = 3). Statistical analysis was performed using one-way ANOVA followed by Tukey’s *post-hoc* test. *p\*\** < 0.01, *p\*\*\** < 0.001, ns; not significant.

Conversely, we examined whether activation of lysosomal biogenesis could alleviate Tau aggregate accumulation. The transcription factor EB (TFEB) is a master gene for lysosomal biogenesis and has been targeted for drug discovery in cancer and neurodegenerative disorders, leading to the development of activators and other modulators. (Settembre *et al*. 2011). Curcumin analog C1 specifically binds to TFEB and promotes TFEB nuclear translocation without inhibiting MTOR activity (Song *et al*. 2016). As a result of the uptake analysis using LysoPrime Deep Red and pHLys Green dyes, treatment with curcumin analog C1 was found to enhance lysosomal acidification and abundance in both WT and G3BP1 KO cells (Figure 5A, 5B). Moreover, TFEB activation was quantified using TFEB-mNG as the ratio of nuclear TFEB-mNG fluorescence intensity to total cellular TFEB-mNG fluorescence intensity. Similar to rapamycin treatment, curcumin analog C1 treatment significantly increased the nuclear translocation of TFEB-mNG (Figure 5C, 5D). After pretreatment with curcumin analog C1 for 1 day, cells were seeded with PFFs, and the accumulation of TauRD-GFP aggregates was assessed 2 days later (Figure 5E). In G3BP1 KO cells, the accumulation was reduced to levels comparable to those observed in untreated WT cells. (Figure 5F). This result suggests that the lysosomal activity reduced by G3BP1 deficiency was elevated by curcumin analog C1 treatment, thereby promoting the degradation of TauRD-GFP aggregates.

**Figure 5.**
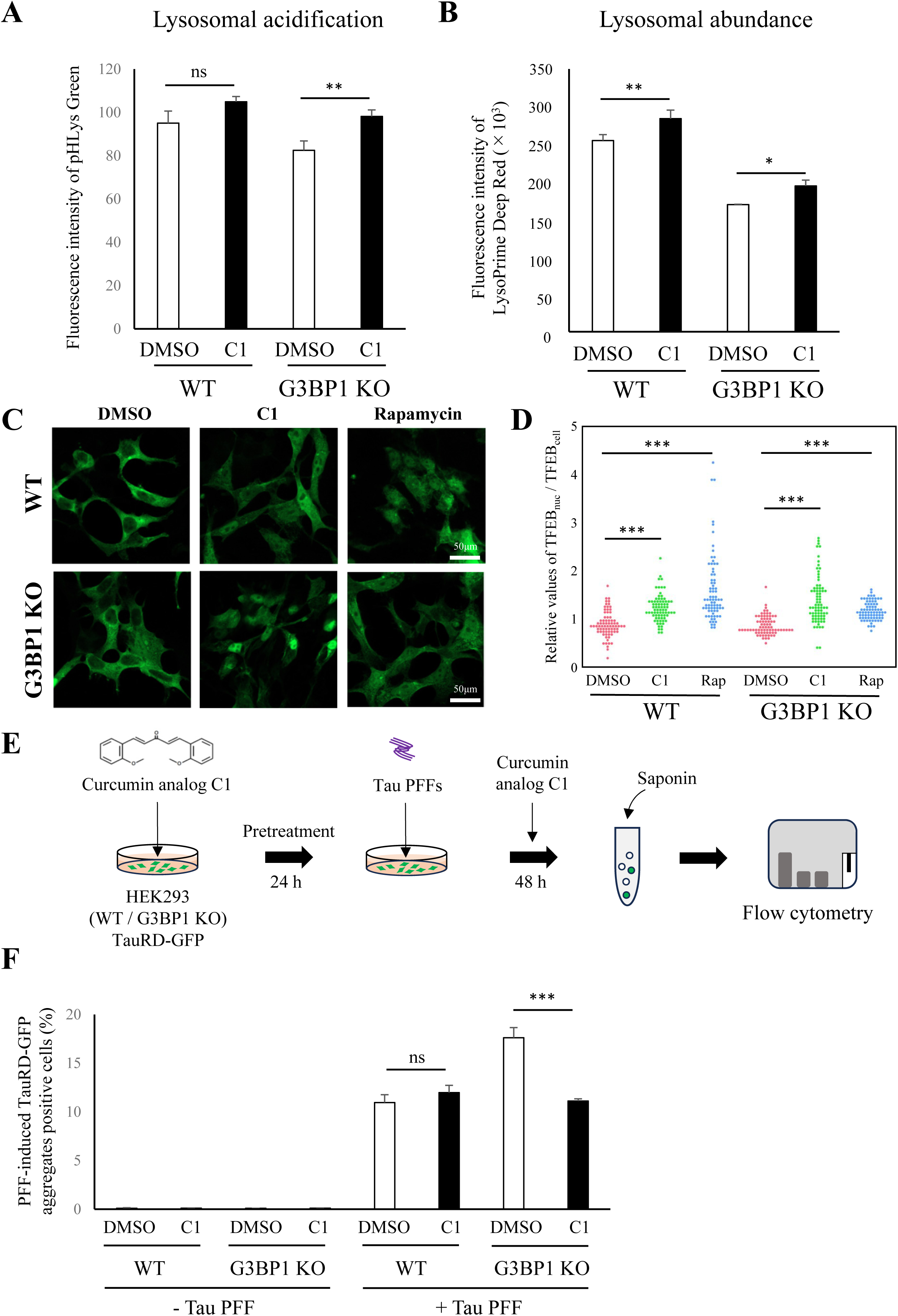
Attenuation of PFF seed-induced Tau aggregation by TFEB activator curcumin analog C1 in G3BP1 KO cells. **A.** Lysosomal acidification was assessed by measuring the uptake of the fluorescent dye pHLys Green in cells treated with the TFEB activator curcumin analog C1. The data are presented as means ± SD (*n* = 3). Statistical analysis was performed using Student’s *t*-test. *p\*\** < 0.01, ns; not significant. **B**. Similarly, lysosomal abundance was measured using LysoPrime Deep Red. The data are presented as means ± SD (*n* = 3). Statistical analysis was performed using Student’s *t*-test. *p\** < 0.05, *p\*\** < 0.01. **C**. To measure TFEB activity, TFEB-mNG was expressed in WT and G3BP1 KO cells. Following treatment with curcumin analog C1 or rapamycin, cells were imaged using confocal laser scanning microscopy. **D**. The nuclear and total cellular TFEB-mNG fluorescence intensities were quantified, and the ratio of nuclear (TFEB_nuc_) to total cellular TFEB-Mng (TFEB_cell_) was plotted. A total of 76 cells were analyzed for each group. Statistical significance was determined by two-way ANOVA followed by Tukey’s multiple-comparison test. Two-way ANOVA revealed significant effects of treatment (F_2,450_ = 72.72, *p* < 0.0001) and genotype (F_1,450_ = 6.74, *p* = 0.0097), as well as a significant treatment × genotype interaction (F_2,450_ = 23.11, *p* < 0.0001), indicating that the effects of curcumin analog C1 and rapamycin on TFEB nuclear translocation. *p\*\*\** < 0.001. **E**. Cells expressing TauRD-GFP were pretreated with the TFEB activator curcumin analog C1 for 24 h and then seeded with PFFs. Cells were subsequently treated with curcumin analog C1 for 48 h, followed by assessment of TauRD-GFP aggregation. **F**. Results of the TauRD-GFP aggregation assay in WT and G3BP1 KO cells are shown. Open bars (DMSO) represent DMSO-treated cells, and closed bars (C1) represent cells treated with curcumin analog C1. The data are expressed as means ± SD (*n* = 3). Statistical analysis was performed using one-way ANOVA followed by Tukey’s *post-hoc* test. *p\*\*\** < 0.001, ns; not significant.

## 4 DISCUSSION

In this study, we performed a pooled CRISPR library screen to identify genes involved in the accumulation of Tau aggregates. The candidate genes were significantly enriched in factors related to mRNA metabolic processes and autophagy. Given that autophagy is a principal protein degradation system, impairment of this pathway can be expected to exert a profound impact on the clearance of Tau aggregates. Indeed, it has been reported that reduced autophagic activity leads to the accumulation of pathological Tau, and that autophagosome accumulation is observed in the brains of patients with tauopathies (Piras *et al*. 2016; Jia *et al*. 2025). On the other hand, the association between mRNA metabolic processes and neurodegenerative diseases has recently attracted considerable attention. RNA transcription, processing, degradation, and stress granule formation are all integral components of mRNA metabolic processes. Previous genetic studies of familial ALS have identified several RNA-binding proteins, such as TDP-43, FUS, and TIA-1, highlighting the importance of RNA metabolism in disease pathogenesis (Mann & Donnelly 2021). These factors play roles in splicing regulation and in the assembly and dynamics of stress granules (Mann & Donnelly 2021; Brown *et al*. 2022). Interestingly, *in vitro* experiments have shown that Tau also binds to RNA, undergoes condensation, and converts into pathogenic Tau with seeding activity (Hochmair *et al*. 2022). Furthermore, RNA-binding proteins containing polyserine stretches, such as SRRM2 and PNN, have been shown to act as nucleation sites of aggregation, thereby facilitating the propagation of Tau aggregates (Lester *et al*. 2023). These results collectively imply that a large proportion of the factors contributing to Tau aggregate accumulation are associated with autophagy and mRNA metabolism.

G3BP1 is an RNA-binding protein known to be involved in stress granule formation. Stress granules are dynamic, non-membranous cytoplasmic assemblies of RNA and proteins that form in response to various cellular stresses, such as oxidative stress, heat shock, or viral infection, and they play important roles in translational regulation, cell protection and RNA and protein quality control (Protter & Parker 2016). While loss of G3BP1 by itself has little effect on stress granule formation, combined loss of G3BP1 and its paralog G3BP2 prevents their assembly. Therefore, the accumulation of Tau aggregates caused by G3BP1 deficiency in this study is thought to be unrelated to defects in stress granule formation. Moreover, it has recently been reported that Tau directly binds to G3BP2 to suppress aggregation, whereas it does not bind to G3BP1 (Wang *et al*. 2023). These findings suggest that G3BP1 is involved in the accumulation of Tau aggregates through a distinct mechanism. Recently, G3BP1 has also been reported to localize to the cytoplasmic surface of lysosomes, where it suppresses mTORC1 signaling through the TSC complex and protects damaged lysosomes, thereby contributing to the regulation of lysosomal function (Prentzell *et al*. 2021; Jia *et al*. 2022). In this study, we measured lysosomal activity in G3BP1-deficient cells. As a result, a significant decrease in lysosomal abundance and an increase in lysosomal acidification were observed in the KO cells, suggesting that the loss of G3BP1 results in impaired lysosomal function. This lysosomal dysfunction could potentially delay the clearance of Tau aggregates. Another research group has reported that knockdown of G3BP1 results in an increase in mutant huntingtin protein (HTT) (Gutiérrez-Garcia *et al*. 2023). A possible involvement of proteasomal degradation mediated by the interaction between mutant HTT and G3BP1 has been suggested, although the underlying mechanisms remain unclear. While we did not investigate proteasome activity in this study, the lack of direct binding between G3BP1 and Tau implies that, unlike in the case of mutant HTT, the proteasome is less likely to be involved in Tau aggregate accumulation.

Lysosomal dysfunction has been implicated as a cause of various neurodegenerative diseases, including AD, PD and lysosomal storage diseases (Zhang *et al*. 2025; Root *et al*. 2021). A variety of pathogenic proteins, such as α-synuclein, and amyloid precursor protein undergo clearance in lysosomes via autophagy and endosomal trafficking. α-Synuclein can be degraded by multiple autophagic pathways, including chaperone-mediated autophagy and macroautophagy (Cuervo *et al*. 2004; Watanabe *et al*. 2012). It has also been suggested that the endosomal trafficking of APP and its C-terminal fragments, which are degraded through the endosome–lysosome pathway, may be regulated by AD-associated factors such as SORL1 and PICALM (Nixon 2017). Conversely, lysosomal damage caused by compounds such as LLOMe has been shown to significantly enhance FPP seed–induced α-synuclein aggregation (Kakuda *et al*. 2024). Similar findings have been reported for Tau, and interestingly, it has been demonstrated that PFF seeds can leak through nanoscale lysosomal damage and act as nucleation sites on the lysosomal membrane surface, thereby initiating the formation of Tau aggregates (Rose *et al*. 2024; Chen *et al*. 2019). Moreover, nanoscale lysosomal lesions are repaired by the ESCRT (endosomal sorting complex required for transport) pathway, and dysfunction of this mechanism results in enhanced leakage of PFF seeds. Thus, the ESCRT-mediated repair of lysosomal membrane damage appears to be critical for limiting Tau aggregate formation (Rose *et al*. 2024). Our findings that G3BP1 deficiency leads to impaired lysosomal activity and enhanced Tau aggregate accumulation may be related to recent reports showing that G3BP1-positive stress granules cover sites of lysosomal damage and facilitate ESCRT-dependent membrane repair (Bussi *et al*. 2023). Taken together, we propose that lysosomal preservation mediated by G3BP1 may play a critical role in the degradation of Tau aggregates. In WT cells, even when lysosomal damage occurs, G3BP1 promotes lysosomal repair and activity, thereby maintaining lysosomal homeostasis and facilitating the degradation of Tau aggregates. In contrast, in G3BP1-deficient cells, both lysosomal repair and activity are impaired even under basal conditions, leading to the accumulation of Tau aggregates. Interestingly, Tau aggregates accumulated to a greater extent in G3BP1-KO cells under basal conditions than in WT cells exposed to LLOMe (Figure 4). This raises the possibility that lysosomal impairment alone does not fully account for the phenotype of G3BP1-KO cells.

The reduction in lysosomal abundance and acidification caused by G3BP1 deficiency was rescued by activation of TFEB (Figure 5A, 5B). As a result of this effect, the accumulation of Tau aggregates was suppressed in G3BP1 KO cells. In contrast, although TFEB activator treatment increased lysosomal abundance and acidification in WT cells, it had no impact on Tau aggregate accumulation, implying that lysosomal activity in WT cells may already be sufficient for Tau clearance. Based on these findings, when lysosomal homeostasis is impaired by aging or environmental factors, therapeutic approaches targeting TFEB-mediated lysosomal biogenesis may prove effective in tauopathies. TFEB-mediated activation of lysosomal biogenesis has long been considered a promising therapeutic target for many neurodegenerative diseases. The curcumin analog C1 used in this study has been reported to activate lysosomal biogenesis in the rat brain following oral administration (Song *et al*. 2016). Indeed, oral administration of curcumin analog C1 ameliorated both APP and Tau pathology in 3xTg-AD mice, one of the most used transgenic models of AD (Song *et al*. 2020). Taken together with these findings, this compound represents a promising candidate for the treatment of tauopathies.

In addition to this compound, other agents that activate lysosomal function, including lithium and AZP2006, have been investigated, and some have progressed into clinical trials for neurodegenerative diseases. Lithium, clinically used as a mood stabilizer for bipolar disorder, has been shown to enhance lysosomal function through both the inositol depletion pathway via inhibition of inositol monophosphatase and the activation of TFEB mediated by GSK3β inhibition (Sarkar *et al*. 2005). Recent study has demonstrated that lithium levels are reduced in the brains of patients with mild cognitive impairment and AD. Moreover, supplementation with lithium orotate has been shown to prevent or ameliorate amyloid-β deposition, Tau phosphorylation, and cognitive decline in mouse models (Aron *et al*. 2025). AZP2006 is a compound that activates lysosomal function through a distinct mechanism from TFEB activation. Specifically, it enhances the trafficking of the Progranulin–Prosaposin complex, thereby promoting lysosomal homeostasis (Aron *et al*. 2025). For this compound as well, safety has been confirmed in humans, and clinical trials are currently underway for the treatment of AD and PSP (Verwaerde *et al*. 2024). Taken together with previous findings, our findings suggest that stabilization of lysosomal homeostasis by G3BP1 may contribute to the prevention or treatment of tauopathies. Although further studies will be required to clarify the underlying mechanisms and translational relevance, G3BP1 itself could potentially serve as a novel therapeutic target in the future.

## ACKNOWLEDGMENTS

We thank Professor Noboru Mizushima (The University of Tokyo, Graduate School and Faculty of Medicine, Tokyo, Japan) for the kind gift of the GFPLLC3LRFP plasmid. This work was supported by Grants-in-Aid for Scientific Research from the Japan Society for the Promotion of Science (21K07422, 24K10494 to Y.W.), AMED under Grant Number JP24wm0625501 (A.H.), and JST SPRING, Japan Grant Number JPMJSP2165 (T.K.).

## DATA AVAILABILITY STATEMENT

The data that support the findings of this study are available upon request.

## AUTHOR CONTRIBUTIONS

**Tomo Kimura**: formal analysis, investigation, validation, visualization, original draft preparation. **Riku Saito**: investigation, validation. **Mariko Okada**: investigation, validation. **Satoaki Matoba**: review & editing. **Atsushi Hoshino**: conceptualization, funding acquisition, investigation, resources, validation, review & editing. **Yoshihisa Watanabe**: conceptualization, data curation, funding acquisition, investigation, project administration, resources, supervision, validation, visualization, original draft preparation, review & editing.

## Abbreviations

AD: Alzheimer’s disease
FTD: frontotemporal dementia
PSP: progressive supranuclear palsy
CBD: corticobasal degeneration
NFTs: neurofibrillary tangles
PD: Parkinson’s disease
ALS: amyotrophic lateral sclerosis
PFF: preformed fibril
LLOMe: L-Leucyl-L-Leucine methyl ester
WT: wild-type
TFEB: transcription factor EB
HTT: huntingtin protein
ESCRT: endosomal sorting complex required for transport

