## Supplemental Figures for "G3BP1 Maintains Lysosomal Homeostasis to Limit Tau Aggregate Accumulation"

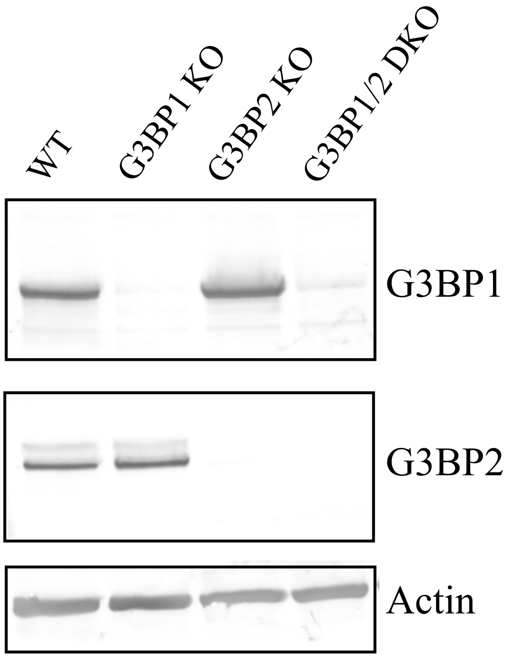


**Figure S1 Western blotting analysis of G3BP1 and G3BP2 KO cells**
To confirm gene disruption in CRISPR–Cas9–edited clones, cell lysates were analyzed by western blotting with anti-G3BP1 (*upper*), anti-G3BP2 (*middle*) or anti-Actin (*lower*) antibodies. *Lane 1*, WT; *lane 2*, G3BP1KO; *lane3*, G3BP2 KO; *lane 4*, G3BP1/2 DKO.

**Figure S1**

**A**

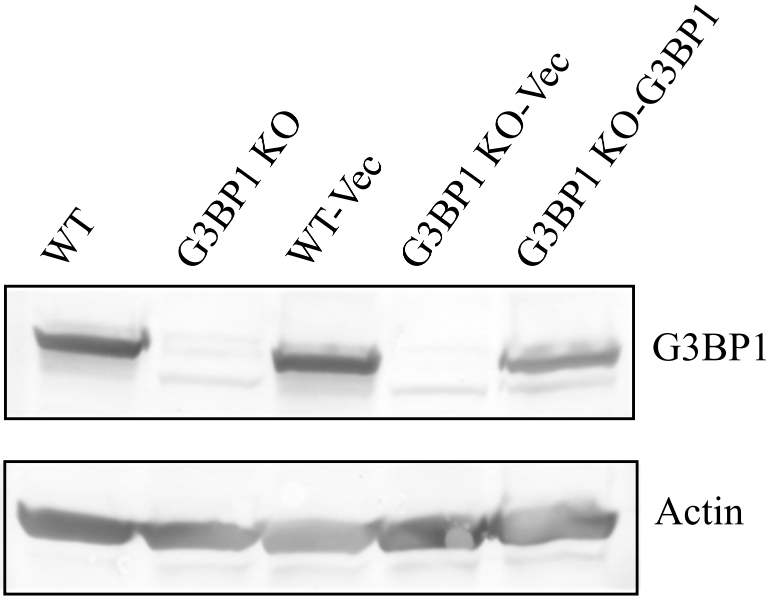


**B**

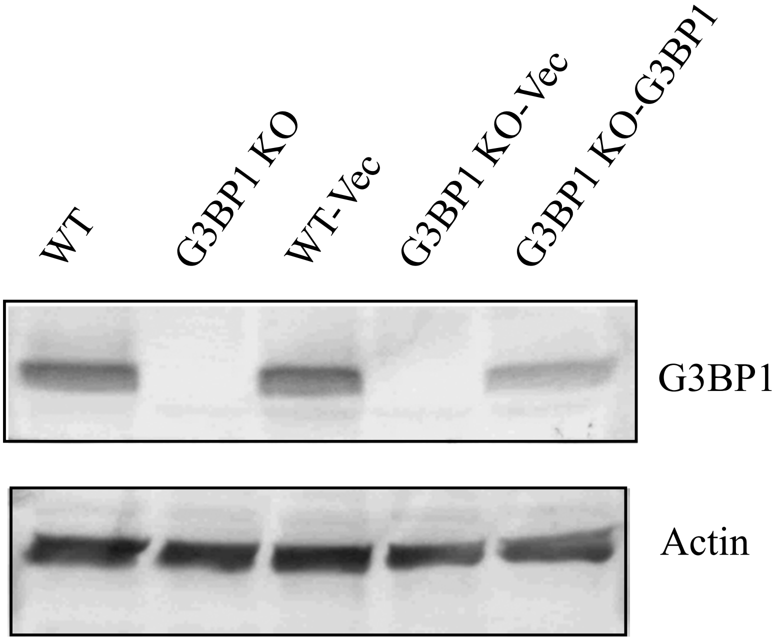


**Figure S2 Western blotting analysis of exogenous expression of G3BP1**

**A**. Exogenous expression of G3BP1 in Fig. 2B was confirmed by western blotting with anti-G3BP1 (*upper*) or anti-Actin (*lower*) antibodies. **B**. Exogenous expression of G3BP1 in Fig. 3B was confirmed by western blotting with anti-G3BP1 (*upper*) or anti-Actin (*lower*) antibodies.

**Figure S2**

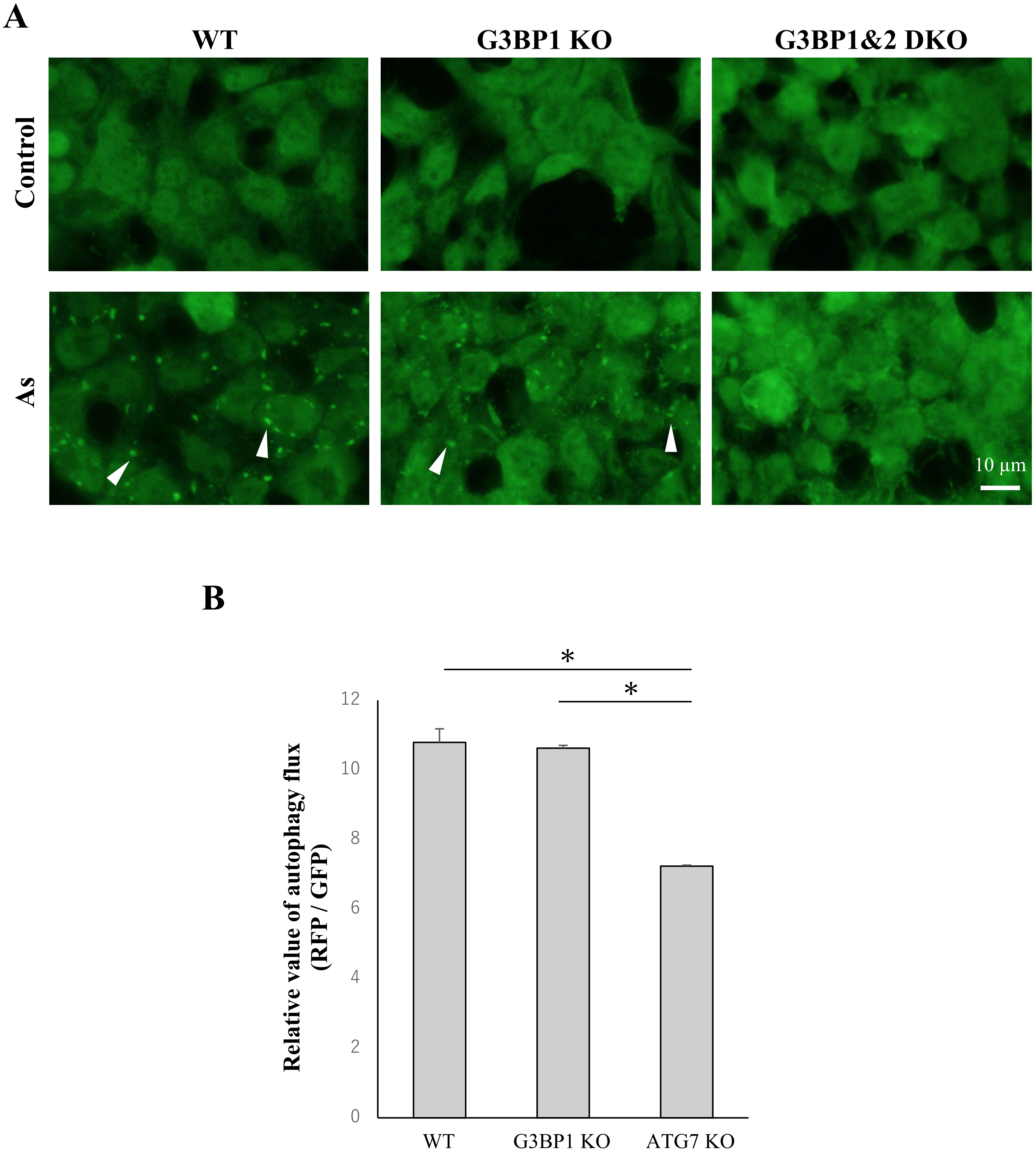


**Figure S3 Assessment of stress granule formation and autophagy flux in G3BP1 KO cells**

**A**. To induce oxidative stress, cells were treated with sodium arsenite, and stress granules were visualized by immunostaining with the stress granule marker TIA-1. Images were acquired by confocal laser microscopy. The panels show cells of WT (*left*), G3BP1 KO (*center*), and G3BP1/2 double KO (*right*), respectively. Arrowheads denote stress granules positive for TIA-1. **B**. For the assessment of autophagic flux, cells were generated to stably express GFP-LC3-RFP. Fluorescence intensities of GFP and RFP were quantified by flow cytometry, and the RFP/GFP ratio was calculated to represent relative autophagic activity. The data are expressed as means ± SD ($n$ = 3). Statistical analysis was performed using one-way ANOVA followed by Tukey's *post-hoc* test. $p$*** < 0.001, ns; not significant.

**Figure S3**

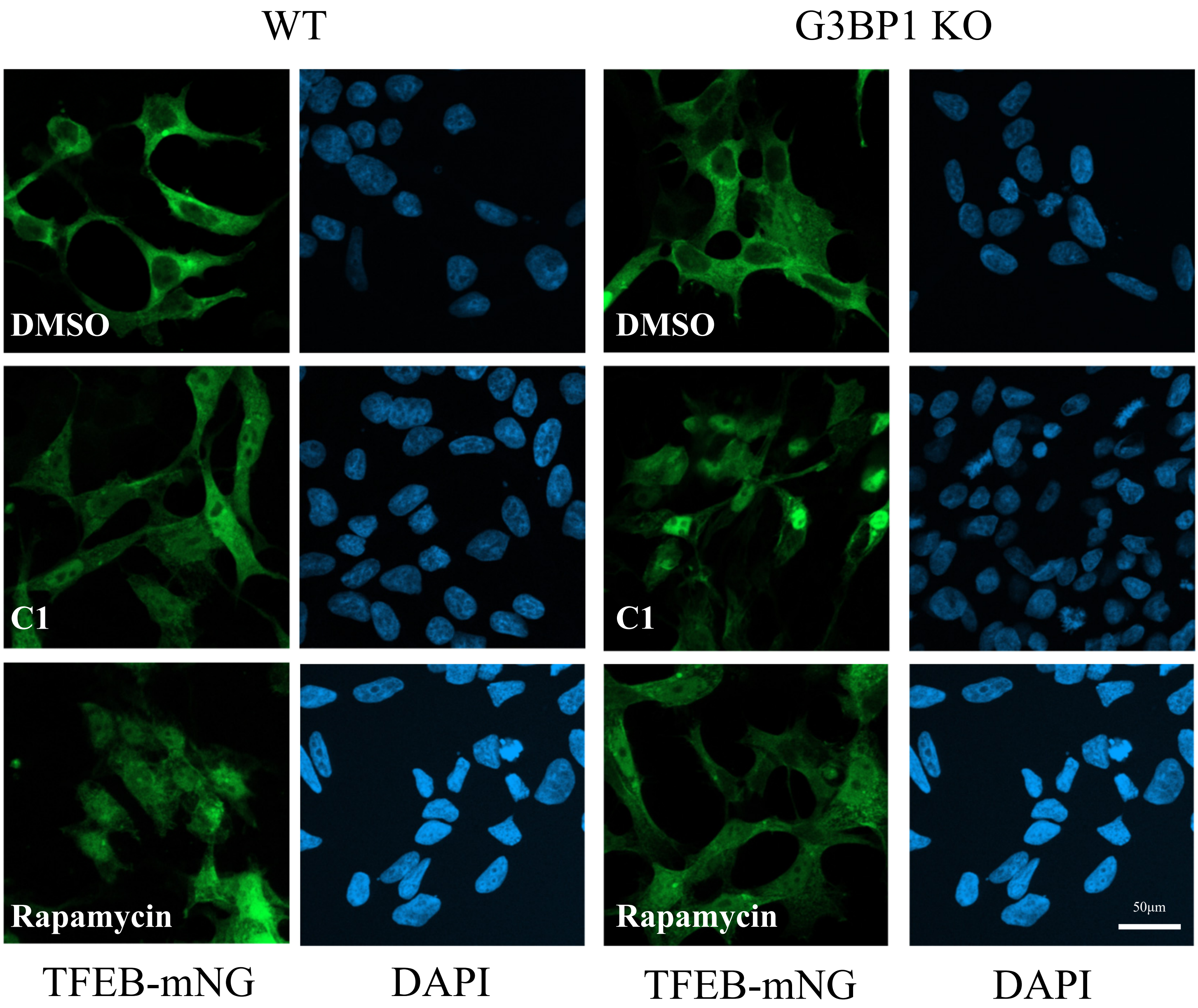


**Figure S4 Nuclear translocation assay of TFEB-mNG**

TFEB activity was assessed using TFEB-mNG. TFEB-mNG was expressed in HEK293 WT and G3BP1 KO cells using a lentiviral vector, followed by treatment with 1 μM curcumin analog C1 or 10 μM rapamycin for 24 h. Nuclei were stained with DAPI, and images were acquired using confocal laser scanning microscopy.

**Figure S4**
